# IFT88 controls NuMA enrichment at k-fibers minus-ends to facilitate their reincorporation into mitotic spindles

**DOI:** 10.1101/512061

**Authors:** Nicolas Taulet, Audrey Douanier, Benjamin Vitre, Christelle Anguille, Justine Maurin, Yann Dromard, Virginie Georget, Benedicte Delaval

**Author notes:** NT and AD have contributed equally to the work.

## Abstract

To build and maintain mitotic spindle architecture, molecular motors exert spatially regulated forces on microtubules (MT) minus-ends. This spatial regulation is required to allow proper chromosomes alignment through the organization of kinetochore fibers (k-fibers). NuMA was recently shown to target dynactin to MT minus-ends and thus to spatially regulate dynein activity. However, given that k-fibers are embedded in the spindle, our understanding of the machinery involved in the targeting of proteins to their minus-ends remains limited. Intraflagellar transport (IFT) proteins were primarily studied for their ciliary roles but they also emerged as key regulators of cell division. Taking advantage of MT laser ablation, we show here that IFT88 concentrates at k-fibers minus-ends and is required for their re-anchoring into spindles by controlling NuMA accumulation. Indeed, IFT88 interacts with NuMA and is required for its enrichment at newly generated k-fibers minus-ends. Combining nocodazole washout experiments and IFT88 depletion, we further show that IFT88 is required for the reorganization of k-fibers into spindles and thus for efficient chromosomes alignment in mitosis. Overall, we propose that IFT88 could serve as a mitotic MT minus-end adaptor to concentrate NuMA at minus-ends thus facilitating k-fibers incorporation into the main spindle.

## INTRODUCTION

The mitotic spindle is a highly dynamic structure that is essential for mammalian cell division^1^. Indeed, proper bipolar mitotic spindle architecture is essential for both spindle orientation and for proper chromosomes alignment in metaphase. To assemble itself and preserve its integrity, the mitotic spindle must continuously coordinate microtubule (MT) nucleation with the integration of various MT structures^1–3^. Indeed, at mitotic entry, peripheral non-centrosome-associated MTs form clusters that move and integrate into the forming spindle^4,5^. Similarly, kinetochore nucleated k-fibers continuously need to be integrated into the main spindle by sliding towards the poles to ensure efficient spindle assembly and proper chromosomes alignment^6–8^. Such a coordination is achieved by the action of MT associated proteins (MAPs) and motors that exert spatially regulated forces on MTs to cluster them into poles. MT minus-ends, which are clustered at the spindle poles, can be considered as a platform for such a regulation. Indeed, NuMA was recently shown to target dynactin to minus-ends and thus spatially regulate dynein activity^8,9^. If proteins recruitment at MT plus-end has been well-documented, the specific targeting of proteins to MTs and k-fiber minus-ends has yet to be fully understood^10^. Indeed, continuous reintegration of k-fibers happens inside an established spindle and is essential to preserve its integrity and to ensure proper chromosomes alignment^6–8^. However, given that k-fibers are embedded in the main spindle, our understanding of the machinery involved in the targeting of proteins to their minus-ends remains limited.

The intraflagellar transport (IFT) machinery is a well conserved intra-cellular transport system that has been studied for a long time for its role in cilia formation and function in non-dividing cells^11^. During IFT-mediated transport, kinesin and dynein motors drive the bidirectional transport of IFT trains from the base to the tip of the cilium in an anterograde movement and from the tip to the base in a retrograde movement^12^. IFTs were therefore accepted to function as cargos, for example, of axoneme precursors such as tubulins as well as molecules of the signal transduction machinery inside the cilium^13,14^. More recently, IFTs were shown to contribute to ciliary motor activation^15^ indicating that a lot remains to be done to fully understand the roles of this complex machinery. Interestingly, in addition to their ciliary roles, IFTs were also shown to function in interphase in the regulation of MT dynamics in the cytoplasm^16^. Moreover, they were shown to contribute to intracellular transport in non-ciliary systems such as lymphocytes^17,18^ or dividing cells^19,20^. Indeed, in mitosis, IFT88 was previously shown to function as part of a dynein1-driven complex required for the transport of peripheral MT clusters to spindle poles to ensure proper formation of astral MT arrays and correct spindle orientation^19^. Recent results also indicate that IFT88 is required for central spindle organization^20^. However, the requirement of IFT proteins for k-fibers organization has never been directly addressed.

Here, taking advantage of MT laser ablation^8^, we show that IFT88, a core member of the IFT machinery, concentrates at k-fibers minus-ends and is required for their re-anchoring into spindles by controlling NuMA accumulation. Mechanistically, IFT88 interacts with NuMA and is required for its enrichment at newly generated k-fibers minus-ends. Combining nocodazole washout experiments and IFT88 depletion, we further confirm that IFT88 is required for the reorganization of k-fibers into spindles and subsequent efficient chromosomes alignment in mitosis. These findings identify a new mechanism for NuMA enrichment at k-fibers minus-ends involving a core member of the IFT machinery. Indeed, we propose that IFT88 could serve as a mitotic minus-ends adaptor to concentrate NuMA at k-fiber minus-ends thus facilitating their incorporation into the main spindle and subsequent chromosomes alignment.

## RESULTS

### IFT88 concentrates at the newly generated k-fiber minus-end after laser ablation

The spindle must continuously coordinate MTs nucleation with the integration of kinetochores bound k-fibers in order to properly control chromosomes alignment. Poleward movement of k-fibers thus continuously happens inside an established spindle^8^ but is hard to observe due to the dense array of MTs in this region. Indeed, k-fibers are embedded in the spindle and their minus-ends are difficult to image. To overcome this challenge and test whether IFT88 could be required for k-fibers reintegration into spindles, we took advantage of MT laser ablation to create new and isolated k-fibers minus-ends in an established spindle. This technique allows to visualize in bipolar or monopolar spindles the reintegration and subsequent transport of both MTs and k-fibers into the spindle^8^. Using immunofluorescence staining, we first found that, after k-fiber laser ablation, IFT88 localizes at newly generated minus-ends (Fig. 1a). To dynamically visualize protein recruitment at k-fibers minus-ends after laser ablation and determine if IFT88 could be recruited at this site, we then performed time-lapse imaging on an Emerald-IFT88 cell line. Upon laser ablation, Emerald-IFT88 was found to rapidly concentrate at the newly generated MT minus-end before decreasing as fibers re-anchor to the main spindle (Fig. 1b and c; Supplementary movie 1). This accumulation of IFT88 at the newly generated minus-end was consistent with the idea that IFT88 could be required for k-fibers reintegration into spindles.

**Figure 1:**
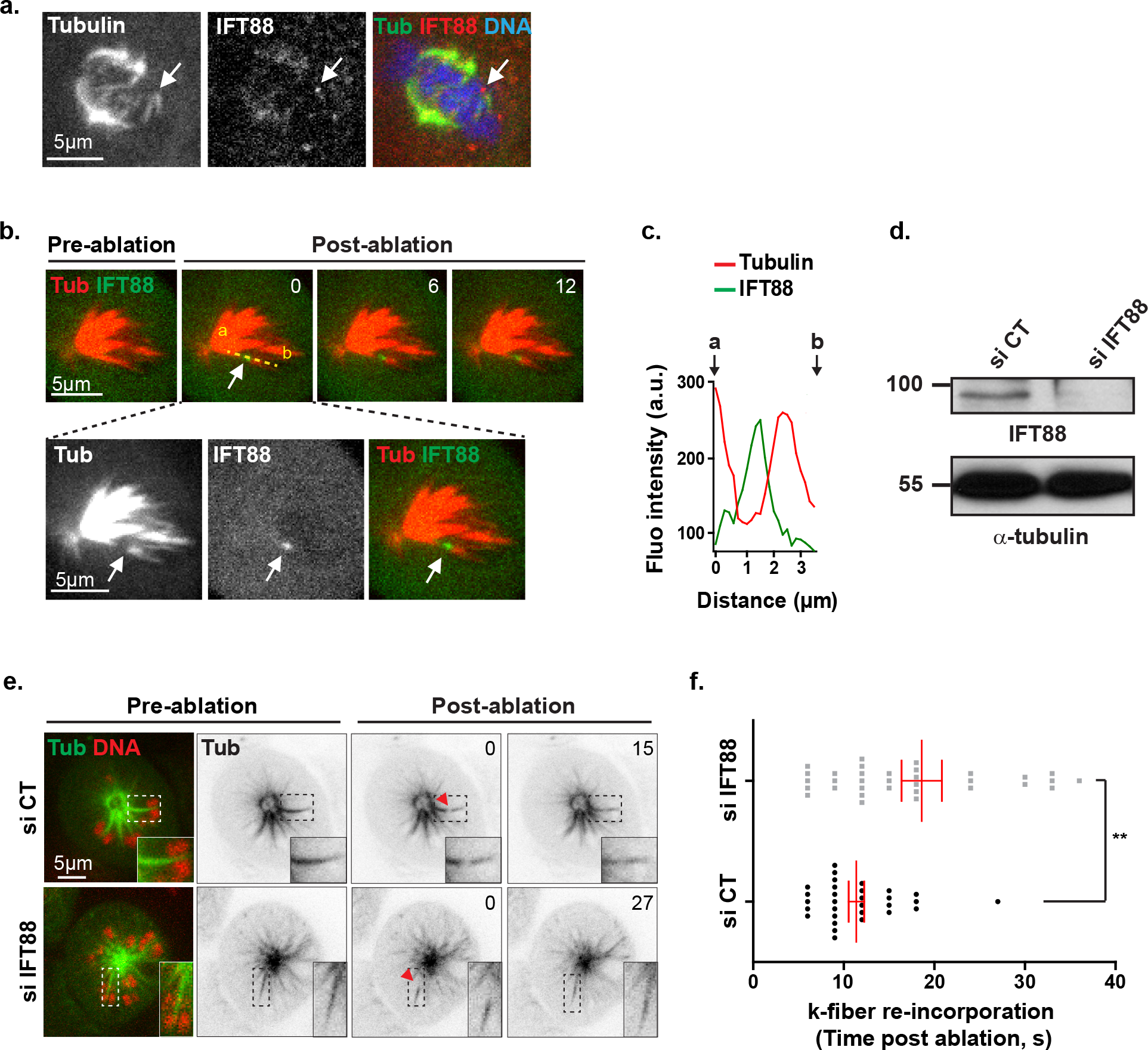
IFT88 is recruited at k-fibers minus-ends after laser ablation and contributes to their re-anchoring into spindle. **a.** Immunofluorescence images of GFP-α-tubulin LLC-PK1 cell (maximal intensity projection) after k-fiber laser ablation in a bipolar spindle showing IFT88 accumulation at minus-end (white arrow). α-tubulin, IFT88 and α-tubulin/IFT88/DNA stainings are shown. **b.** Images from time-lapse microscopy of monopolar Emerald-IFT88 LLC-PK1 cells labelled for tubulin (SiR-Tubulin) before and after k-fiber laser ablation (upper panel). Hoechst live was used to identify k-fibers attached to chromosomes. Time post-ablation (s). α-tubulin, IFT88 and α-tubulin/IFT88 stainings (maximal intensity projection of 2 planes) show IFT88 accumulation at minus-end after laser ablation (white arrow). **c.** Line scans representing α-tubulin and IFT88 fluorescence intensities measured from a to b along the yellow line shown in (**b**). **d.** Western-blots showing the amount of IFT88 in GFP-α-tubulin LLC-PK1 cells transfected with control (CT) or IFT88 siRNA. α-tubulin: loading control. **e.** Images from time-lapse microscopy of monopolar GFP-α-tubulin LLC-PK1 labelled for DNA (Hoechst live, red), to allow for k-fibers detection, in control (CT) and IFT88-depleted cells (left panels). Inverted contrast images of α-tubulin before and after k-fiber ablation (ablation site, red arrowhead) show a delay in k-fiber re-anchoring into spindle upon IFT88 ablation. Time post-ablation (s). Single planes are shown. Insets: magnification of the ablated k-fibers, dashed boxes regions. **f.** Quantification of the time (s) required for k-fiber re-incorporation into spindle after laser ablation in CT and IFT88-depleted cells. n ≥30 ablated k-fibers (1 ablated k-fiber per cell), 3 experiments. Mean +/− s.e.m \*\**P* < 0.01 compared to control (*t* test). Scale bars: 5μm.

### IFT88 contributes to k-fibers re-anchoring into spindles after laser ablation

To test whether IFT88 could participate to k-fibers reincorporation into spindles, we performed time-lapse imaging on GFP-α-tubulin cells^5^ and monitored k-fibers re-anchoring after ablation upon control and IFT88-depleted conditions (Fig. 1d-f; Supplementary movie 2). IFT88 depletion was controlled by western-blot (Fig. 1d) and Hoechst live staining was combined to GFP-α-tubulin to allow chromosomes visualization. In control condition, as previously shown^8^, detached k-fibers rapidly re-anchored to the neighboring MTs and were transported back towards the pole (Fig. 1e and f). In contrast, k-fibers re-anchoring was significantly delayed upon IFT88 depletion (Fig. 1e and f). These results indicate that, upon laser ablation, IFT88, which accumulates at the newly generated minus-end, is required for efficient k-fibers re-anchoring into spindles.

### IFT88 interacts with NuMA and is required for its minus-end enrichment

K-fibers reintegration was previously shown to depend on NuMA accumulation at the newly generated minus-end^8^. Subsequently, dynein activity is recruited at this location to allow proper re-anchoring of ablated MTs to the spindle^9^. However, how NuMA concentrates at k-fiber minus-ends is not completely understood. The rapid enrichment in IFT88 observed at minus-end after ablation (Fig. 1b) suggested that IFT88 could be required for NuMA accumulation at this site. To test this hypothesis, we generated new and resolvable k-fibers minus-ends using laser ablation on monopolar spindles and monitored NuMA enrichment upon control and IFT88-depleted conditions (Fig. 2a-c; Supplementary movie 3). More specifically, YFP-NuMA cell line was used to dynamically monitor by live imaging the impact of IFT88 depletion on the recruitment of NuMA at the newly generated k-fiber minus-end after ablation. As previously described, NuMA rapidly localized at new minus-end structures and accumulated there before their reincorporation into the main spindle (Fig. 2a-c). However, the striking accumulation of NuMA observed at the ablated k-fibers minus-ends in control cells was decreased in IFT88-depleted cells (Fig. 2a-c). Indeed, only 24% of newly generated minus-ends presented with NuMA accumulation compared to 68% in control condition (Fig. 2b). The decrease in NuMA intensity was further validated by fluorescence intensity quantifications (Fig. 2c). To confirm whether NuMA and IFT88 could interact in mitosis, we then performed co-immunoprecipitation (Fig. 2d). HeLa cells expressing Flag-IFT27 and YFP-NuMA were synchronized in mitosis. Flag IP was used, as previously described^20^, to pull-down IFT proteins including IFT88, and revealed an interaction with YFP-NuMA in mitotic cells. These experiments showed that the IFT machinery, including IFT88 and IFT27 can interact with NuMA in mitotic cells and that IFT88 is required for NuMA enrichment at k-fibers minus-ends after ablation. Taken together, these results show that IFT88 contributes to k-fibers re-anchoring into spindles by concentrating NuMA at the new k-fibers minus-ends generated after ablation.

**Figure 2:**
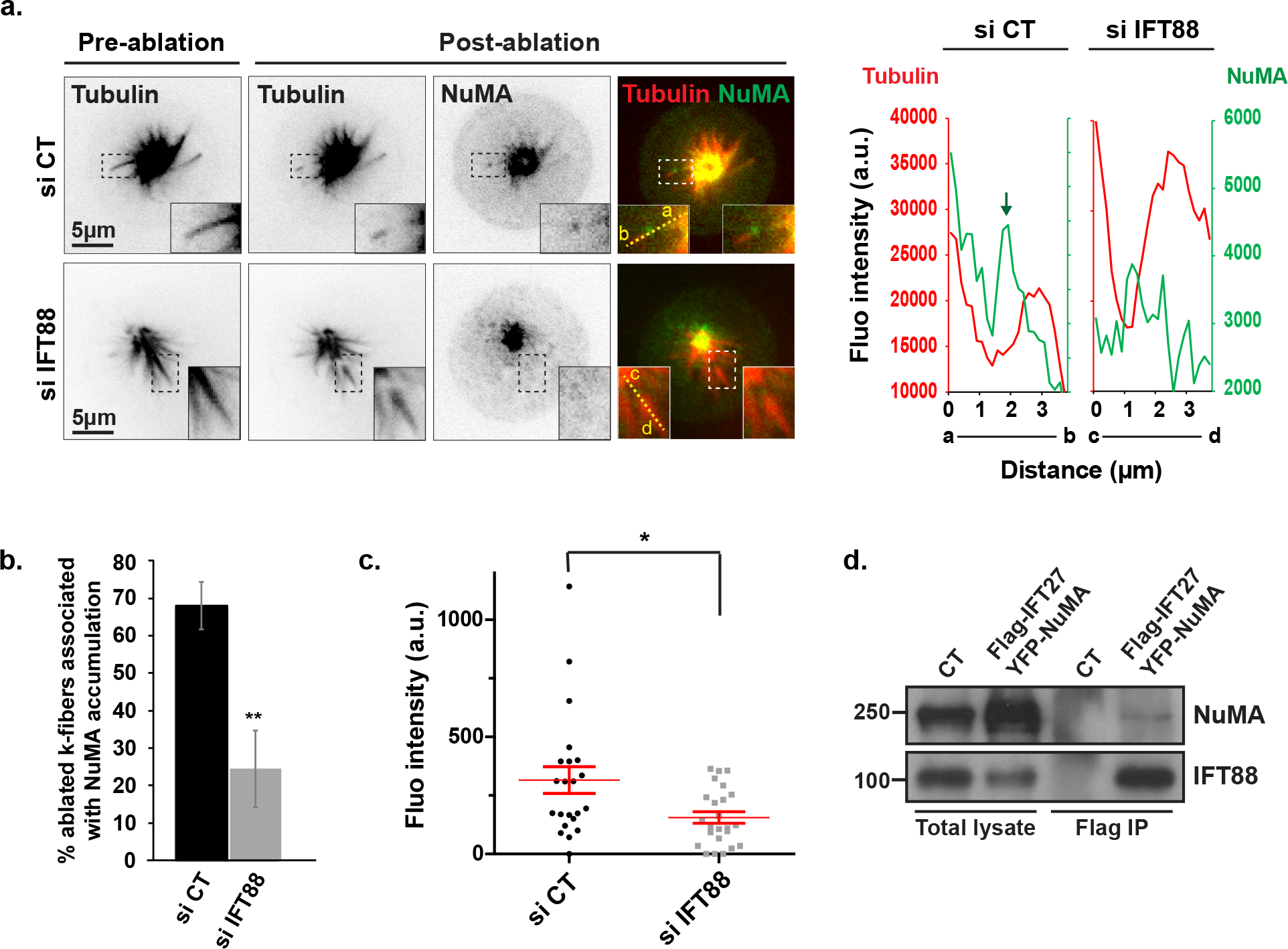
IFT88 interacts with NuMA and contributes to its enrichment at k-fibers minus-ends after laser ablation. **a.** Inverted contrast and merged images from time-lapse microscopy of monopolar YFP-NuMA LLC-PK1 combined with MT labelling (SiR-Tubulin) in control (CT) and IFT88-depleted cells before and after k-fiber laser ablation. The 3s time-point after ablation is shown. Single planes are shown. Insets: magnification of the ablated k-fibers, dashed boxes regions. Hoechst live was used to identify k-fibers attached to chromosomes. Linescans (right) representing NuMA and α-tubulin fluorescence intensities, measured from a to b (control) or from c to d (siRNA IFT88) along the yellow line (left inset on the image), show an accumulation of NuMA at minus-ends of k-fibers after laser ablation in CT cells but not in IFT88-depleted cells. Scale bars: 5 μm. **b.** Percentage of cells with ablated k-fibers associated with NuMA enrichment. n ≥29 ablated k-fibers (1 ablated k-fiber per cell), 3 experiments. Mean +/− s.d. \*\**P* < 0.01 compared to control (*t* test). **c.** Quantification of NuMA fluorescence intensity at minus-ends of MT after laser ablation. n ≥22 cells, 2 experiments. Mean +/− s.e.m. \**P* < 0.05 compared to control (*t* test). **d.** Flag immunoprecipitation performed on mitotic HeLa Kyoto cells transfected with Flag-IFT27 and YFP-NuMA indicates an interaction between YFP-NuMA and IFT proteins, including IFT88. Scale bars: 5 μm.

### IFT88 is required for k-fibers reintegration into spindles upon nocodazole washout and for subsequent chromosomes alignment

As an alternative approach to monitor k-fibers reintegration into spindles, we decided to challenge the spindle using nocodazole to depolymerize MTs and monitor spindle reorganization and chromosomes alignment after washout (Fig. 3a-c; Supplementary movie 4). Indeed, nocodazole washout gives cells the opportunity to nucleate new sets of both centrosomal and acentrosomal MTs, the latest being nucleated in the chromosomal region. These acentrosomal MTs then need to reorganize into k-fibers that rapidly get reincorporated into the spindle to allow for proper chromosomes alignment. Nocodazole washout therefore gives a unique opportunity to monitor the reorganization and reincorporation of k-fibers attached to chromosomes into spindles^6,7,21^. In agreement with a role of IFT88 in k-fibers reorganization after nocodazole washout, acentrosomal MTs nucleated in the chromosomal region did not get properly reincorporated into the main spindle upon IFT88 depletion (Fig. 3a). Indeed, they were delayed in their reincorporation as around 80% of cells still presented with disorganized spindles 5 min after washout compared to 40% in controls (Fig. 3a). Importantly, NuMA minus-end localization was affected by IFT88 depletion as diffuse NuMA staining was observed compared to intense and focused NuMA signal observed in the control conditions (Fig. 3b). This result further strengthened the fact that upon nocodazole washout, IFT88 was required for proper NuMA concentration at minus-ends and subsequent k-fibers reintegration into spindles. Importantly, the delay in spindle reorganization was confirmed by live imaging and correlated with defects in the establishment of a proper metaphase plate (Fig. 3c). Indeed, quantifications indicated that if most control cells already presented with a visible metaphase plate 5 min after washout with only 19% still showing major chromosomes misalignment, 47% of IFT88-depleted cells still presented with major chromosomes misalignment (Fig. 3a). Altogether, these observations strengthen the fact that IFT88 is required for the reorganization of k-fibers into spindles, subsequently contributing to proper chromosomes alignment after nocodazole washout. To further validate the requirement of the IFT machinery in chromosomes alignment, we monitored the impact of IFT88 depletion on chromosomes alignment in mitosis. IFT88 depletion, achieved either by siRNA (Fig. 3d) or using a CRISPR-based auxin inducible degron approach (Fig.3e), led to defects in chromosomes alignment compared to control condition, thus demonstrating that IFT88 is indeed required for efficient chromosomes alignment in mitosis.

**Figure 3:**
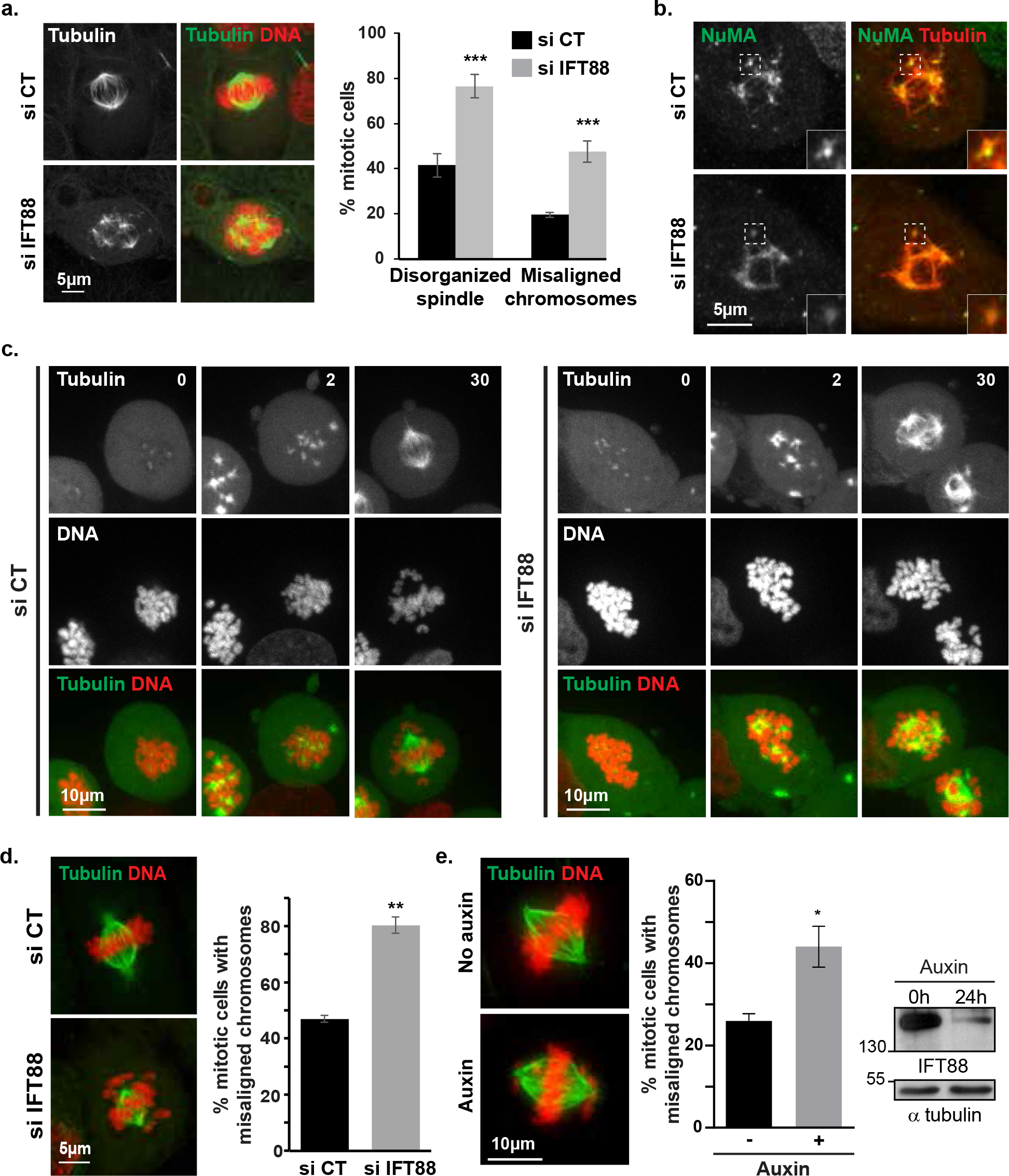
IFT88 contributes to k-fibers reincorporation into spindle after nocodazole washout and is required for proper chromosomes alignment. Immunofluorescence images of GFP-α-tubulin LLC-PK1 upon nocodazole washout showing defects in k-fibers reincorporation into the main spindle in IFT88-depleted cells compared to control. α-tubulin and α-tubulin/DNA stainings are shown (left panel). Percentage of mitotic cells with disorganized spindles or misaligned chromosomes upon nocodazole washout. n >300 mitotic cells. 3 experiments. Mean +/− s.d. \*\*\**P* < 0.001 compared to control (*t* test) (right panel). **b.** Immunofluorescence images of GFP-α-tubulin LLC-PK1 cells upon nocodazole washout showing defects in NuMA minus-ends localization in IFT88-depleted cells. NuMA and α-tubulin/NuMA stainings are shown. Insets: magnified dashed boxes regions. **c.** Images from time-lapse microscopy of LLC-PK1 GFP-α-tubulin/mCherry-H2B cells showing defects in spindle organization and chromosomes alignment in IFT88-depleted cells compared to control upon nocodazole washout. Time after washout (min). **d.** Immunofluorescence images of GFP-α-tubulin LLC-PK1 cells showing defects in chromosomes alignment upon IFT88 depletion. α-tubulin/DNA staining is shown (left). Quantification (right): percentage of mitotic cells with misaligned chromosomes. n > 100 mitotic cells. 3 experiments. Mean +/− s.d. \*\**P* < 0.01 compared to control (*t* test). **e.** Immunofluorescence images (left) showing α-tubulin and DNA stainings in HCT116-AID-IFT88 cells. Control (No auxin) and auxin (30h)-induced AID-YFP-IFT88 degradation conditions are shown. Quantification (right): percentage of mitotic cells with misaligned chromosomes upon auxin treatment (30 h). n >50 mitotic cells. 3 experiments. Mean +/− s.e.m \**P* < 0.05 compared to control (*t* test). Western-blots showing AID-YFP-IFT88 depletion in HCT116 cells upon auxin treatment. α-tubulin: loading control. In all panels, maximum projections are shown, scale bars: 5 or 10 μm.

## DISCUSSION

Collectively, our results identify a novel role for an IFT protein at k-fibers minus-ends in mitosis. Indeed, taking advantage of MT laser ablation, we show that IFT88 rapidly and strongly accumulates at newly generated k-fibers minus-ends upon ablation to efficiently allow their re-anchoring into the main spindle. Moreover, we show here for the first time that IFT88 is required for proper spatial targeting of NuMA at k-fibers minus-ends in mitosis and that IFT proteins, including IFT88, can interact with NuMA in mitotic extracts. This finding is strengthened by nocodazole washout experiments that further confirm the requirement of IFT88 for the reorganization of k-fibers into spindles and thus for efficient chromosomes alignment. Taken together, our results are consistent with a model (Fig. 4) in which, upon perturbations of spindle integrity, IFT88 rapidly identifies k-fibers minus-ends, and is required for an efficient minus-ends re-anchoring response by allowing proper NuMA accumulation. Such an efficient repair mechanism is important to ensure spindle integrity. Indeed, we propose here that IFTs would participate to a pathway that provides both flexibility and robustness for the spindle to adapt to perturbations and ensure efficient chromosomes alignment.

**Figure 4:**
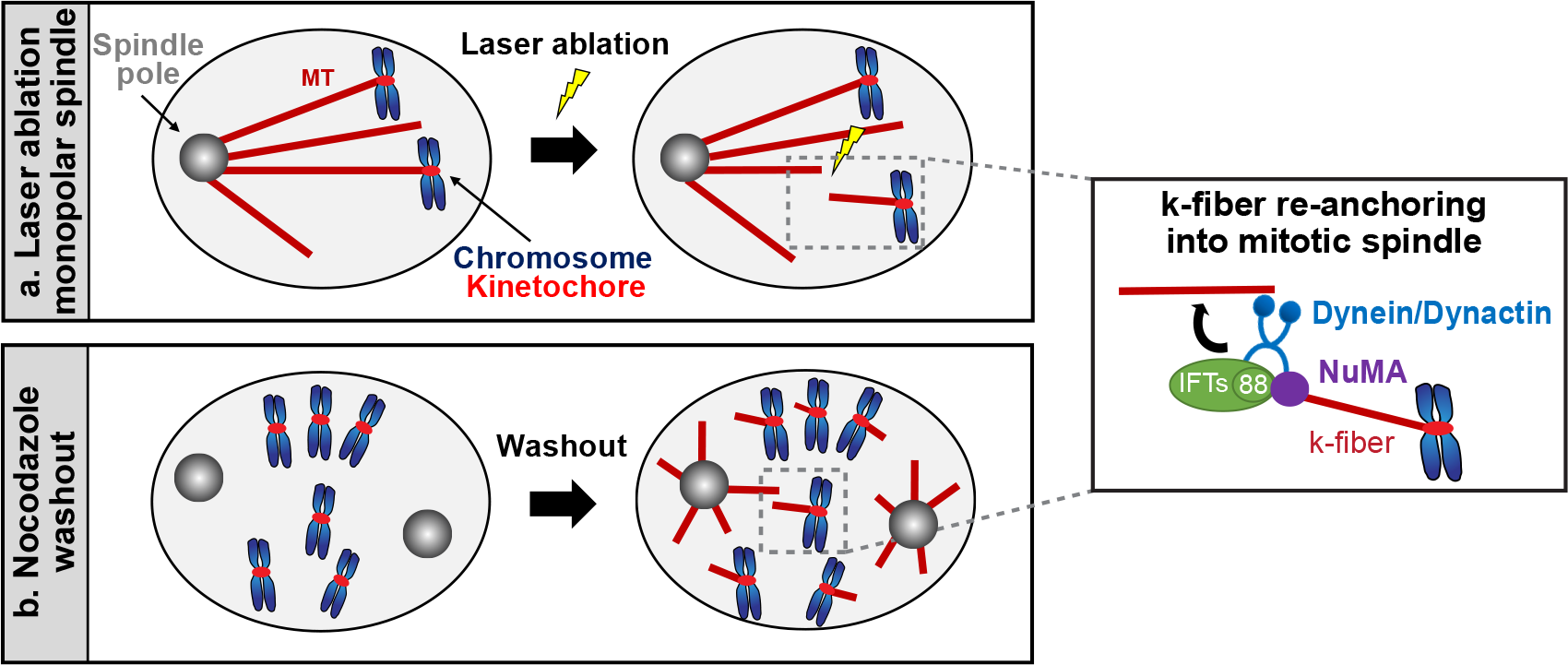
Model for IFT88 function in k-fibers reintegration into mitotic spindles. Upon perturbations, including MT laser ablation (a) or nocodazole washout (b), IFT88 is required for an efficient minus-ends re-anchoring response by allowing proper NuMA accumulation at k-fibers minus-ends. As described previously, NuMA would then be required to target dynein activity to minus-ends through dynactin recruitment. In this model, IFTs, including IFT88, would serve as a mitotic minus-end adaptor to concentrate NuMA at minus-ends thus facilitating k-fibers incorporation into the main spindle.

Our understanding of the mitotic spindle has grown exponentially with the characterization of multiple coexisting MT nucleation pathways that contribute to its assembly, thus ensuring the fidelity of chromosomes segregation^1^. However, we are only starting to understand how all these pathways work together to assemble a functional bipolar spindle and to maintain its integrity. Moreover, all the factors that contribute to the coordination of these different pathways have yet to be fully characterized. NuMA and dynein1 have long been known as factors facilitating spindle assembly and contributing to the maintenance of its integrity^22–26^. Yet, NuMA was only recently shown to provide both spatial and temporal regulation of dynein powered forces^9^. Indeed, MT and k-fibers minus-ends dynamics in mitosis still remain largely less understood than plus-ends dynamics due to the fact that they are embedded into the main spindle and thus difficult to isolate and image. Advanced imaging methods and perturbation tools, including laser ablation, now allow to isolate and track MT minus-ends in space and time^8,27–29^. In this context, our work provides new insights on the regulation of k-fibers dynamics by involving a new player, IFT88, in the control of NuMA local enrichment at minus-ends. By implicating a core member of the IFT machinery in this process, this work strengthens and extends recent works^8,9^ since we propose that IFT88 could serve as a mitotic minus-end adaptor to concentrate NuMA at minus-ends thus facilitating k-fibers reincorporation into the spindle.

This work, together with the previously established roles of IFTs in prometaphase^19^, provides novel insights on how IFT proteins could control spindle mechanical integrity. We previously showed that, upon mitotic entry, IFTs are required for proper spindle assembly and positioning^19^. Indeed, in association with dynein1, they allow for the assembly of robust astral MT arrays by contributing to peripheral MT clusters re-localization towards the poles. These results, combined to earlier works on dynein1 in transporting peripheral MT and centrosome components, demonstrated that IFTs are required to build the mitotic spindle at the entry of mitosis. We show here that, upon perturbations of spindle mechanics, IFTs are required for spindle repair by allowing a rapid enrichment of proteins, such as NuMA which is essential for MT and k-fibers minus-ends re-anchoring to mitotic spindle. Indeed, as described previously^9^, NuMA accumulation at minus-ends is essential to provide the local activation of dynein leading to MT minus-ends transport towards poles. Peripheral MT clusters and k-fibers attached to chromosomes can be considered as ‘pre-assembled’ parts of the spindle, which are reminiscent of ciliary components for which IFTs serve as cargo inside the cilium^13^. Together, these works place IFTs as cargo adaptors to ensure the spatial accumulation of proteins at both MTs and k-fibers minus-ends and to control spindle mechanical integrity, which in turn is essential to ensure proper chromosomes alignment. Whether IFTs only contribute to chromosome alignment by controlling spindle integrity or whether they could also act elsewhere, for example at the kinetochore, will require further investigations. Of note, IFTs also serve as cargo adaptors at later mitotic stages to control central spindle organization^20^.

The mitotic spindle is a highly dynamic structure that requires a robust molecular machinery to maintain its integrity^1,2,30^. If the major pathways for spindle assembly have been extensively characterized, a lot still has to be uncovered on less studied spindle assembly factors that contribute to its integrity, including IFT proteins. This is particularly important for spindle assembly pathways involved in error detection and correction. Indeed, these alternative pathways are crucial to ensure proper chromosomes alignment and segregation and proper completion of mitosis. Future works, including *in vitro* studies to avoid the complexity of the cellular context, will address which IFT proteins participate in the maintenance of spindle mechanical integrity, how IFT sub-complexes interact with MT minus-ends and how their recruitment is regulated. Such experiments will indeed be key to unravel how IFTs, MAPs and motor proteins sequentially get recruited to MT minus-ends and will allow to fully comprehend the contribution of the IFT machinery to spindle mechanical integrity. By characterizing a novel cilia-independent function for an IFT at the minus-ends of k-fibers, this work complements recent evidences that introduce new perspectives on cellular processes involving IFT proteins beyond their role in cilia. These include, in addition to cell cycle^31^ and cell division^19,20,32^, immune synapse polarization^17,18,33^, cell migration^34^ and the regulation of cytoplasmic MT dynamics^16^ or actin^35^. Future works will be required to address whether IFTs function in these non-ciliary roles only as cargo adaptors or also as motor regulators as shown for ciliary motors^15^.

## METHODS

### Cell culture

LLC-PK1 (CLS Cell Lines Services GmbH, Germany), GFP-αtubulin LLC-PK1 (Gift from P. Wadsworth^5^, GFP-αtubulin/mCherry-H2B LLC-PK1 previously generated^20^ and YFP-NuMA LLC-PK1 cells were grown in a 1:1 mixture of Opti-MEM/HAM’s F10 media supplemented with 10% fetal bovine serum (FBS). HCT116-IFT88-AID, previously generated^20^, were grown in DMEM Glutamax medium supplemented with 10% FBS. YFP-NuMA and mEmerald-IFT88 LLC-PK1 cells were generated by transfection of LLC-PK1 with pEYFP-C1-NuMA (Addgene plasmid #28238) or mEmerald-IFT88-N-18 (Addgene plasmid #54125). Cells were then selected with G418 and sorted based on their fluorescence. SiR Tubulin and Hoechst live were added before starting live experiments for 2h at 100 nM and for 30min at 5μg/ml respectively.

### Cell synchronization, inhibitors and microtubules regrowth assays

For immunoprecipitation experiments, HeLa Kyoto cells were synchronized in mitosis using nocodazole 100 ng/ml (Sigma-Aldrich) for 15h followed by 30min release. To induce monopolar spindles, LLC-PK1 cells were treated with the kinesin-5 inhibitor S-trityl-l-cysteine (STLC, Santa Cruz) 2 μM for 15h. For MT regrowth assays, cells transfected for 48h with siRNA were treated with 100 ng/ml nocodazole (Sigma-Aldrich) in culture medium ON at 37°C to depolymerize MTs. Cells were placed on ice for 10 min and washed three times with 4°C PBS. Cells were then incubated in 37°C culture medium without nocodazole and kept at 37°C to allow regrowth. Cells were fixed in MeOH at the indicated time after washout and processed for immunofluorescence. For live imaging, after nocodazole washout in 4°C PBS, cells were incubated in 4°C culture medium instead of 37°C to delay regrowth in order to facilitate imaging.

### AID cell line, siRNA and cDNA transfections

Targeted proteins were depleted with small-interfering RNAs (siRNAs) designed and ordered via Dharmacon (Lafayette, CO), as previously described^20^, and delivered to cells at a final concentration of 100 nM using Oligofectamine (Invitrogen, Carlsbad, CA) according to manufacturers' instructions. The efficacy of proteins knockdown was assessed by immunoblotting 48 h post transfection. HCT116-IFT88-AID cells were generated as previously described^20^. Targeting of IFT88 and degradation of IFT88-AID-YFP was confirmed by western-blot following addition of Auxin (Sigma-Aldrich) at 500 μM in the culture medium for 24h. cDNA transfections were performed using JetPEI (Polyplus Transfection) or Fugene 6 (Promega) transfection reagents according to manufacturers’ instructions for 24h. cDNAs transfected include pCMV14-IFT27-3xFlag^20^, pEYFP-C1-NuMA was a gift from Michael Mancini (Addgene plasmid # 28238; http://n2t.net/addgene:28238; RRID: Addgene_28238) and mEmerald-IFT88-N-18 was a gift from Michael Davidson (Addgene plasmid # 54125; http://n2t.net/addgene:54125; RRID: Addgene_54125).

### Lysates, immunoprecipitations and immunoblotting

HeLa Kyoto, LLC-PK1 and HCT116 cell extracts were obtained after lysis with buffer containing 50 mM Hepes (pH 7.5), 150 mM NaCl, 1.5 mM MgCl_2_, 1 mM EGTA, 1% IGEPAL CA-630 and protease inhibitors (Sigma-Aldrich). Protein concentration for lysates was determined using Bradford reagent (Sigma-Aldrich), loads were adjusted and proteins were resolved by SDS-PAGE and analyzed by western-blot (Western Lightning Plus-ECL kit; PerkinElmer). For Flag immunoprecipitations, HeLa Kyoto cells were transfected with Flag-IFT27 and YFP-NuMA plasmids for 24h and synchronized in mitosis. The resulting cell extracts were incubated for 2h at 4°C with Flag M2 agarose beads (Sigma-Aldrich). Beads were washed four times with 800 μl lysis buffer and the immunoprecipitated proteins were separated by SDS-PAGE and analyzed by western blotting.

### Antibodies

The following primary antibodies were used (western-blot WB, immunofluorescence IF): IFT88 #13967-1-AP (WB: 1/500, IF: 1/250) from Proteintech, α-tubulin (DM1α, Sigma-Aldrich #T6199, WB: 1/400), FITC-conjugated α-tubulin (DM1α, Sigma-Aldrich #F2168, IF: 1/300), NuMA (Santa-Cruz sc-365532 IF: 1:200; Abcam ab-36999 WB:1/500) and DAPI (Cell signaling, IF 1/10000). Secondary antibodies include for IF: Alexa Fluor 488 (#4412S or #4408S) or 555 (#4413S or #4409S) -conjugated anti-rabbit or anti-mouse secondary antibodies (Molecular Probes, 1/1500) and for WB: anti-mouse and anti-rabbit IgG, HRP linked antibody (Cell signaling #7076 and #7074, 1/5000).

### Immunofluorescence

For immunofluorescence experiments, cells were fixed in −20°C MeOH to preserve MTs staining. Then, cells were blocked with PBS-BSA 1%-Triton 0.5% and stained for immunofluorescence with the appropriate primary and secondary antibodies. Slides were mounted in prolong gold (Life Technologies).

### Microscopy, laser ablation and image analysis

Epifluorescence images of the auxin inducible degron experiment were acquired with a Leica DM6000 microscope (Objective: 63×/1.4 NA Plan-Apo) equipped with a Cool SNAP HQ2 camera and controlled by MetaMorph (Molecular Devices). Confocal images and time-lapse were performed using a spinning disk confocal microscope, a Nikon Ti Eclipse coupled to a Yokogawa spinning disk head and an EMCCD iXon Ultra camera (60×/1.4 NA), controlled by the Andor iQ3 software (Andor) or a Zeiss confocal LSM880 (Objective 63×/1.4 NA) controlled by Zen (Zeiss). For laser ablation experiments, the spinning disk confocal microscope (Objective 60×/1.4 NA; optovar 1.5) is coupled to the MicroPoint system (Andor) equipped with a pulsed nitrogen pumped tunable dye laser capable of MT ablation (wavelength 551 nm, frequency 10 Hz, repetition rate 10, number of repeats 5). Three images (plane of the ablation) were acquired at 1 s interval before the ablation, then images were acquired every 3 s for 2 min (5 planes over 4μm; centered on the plane of the ablation). For nocodazole washout experiments, images were acquired every 15s (maximum projections are shown, 11 planes over 10μm). Image processing and analysis (cropping, rotating, brightness, contrast adjustment, color combining and fluorescence intensity measurements (mean intensity of the region of interest minus mean intensity of a background region of the same size inside the cell) were performed with ImageJ. Linescans were obtained using ImageJ plot profile tool. Movies were generated with MetaMorph (Molecular Devices) and displayed at 3 or 10 frame per second as indicated.

### Statistical analysis

The number of cells counted per experiment for statistical analysis is indicated in figure legends. Graphs were created using Microsoft Excel or GraphPad Prism software and error bars represent the s.d. or s.e.m. as indicated. p-values were calculated using a two-tailed Student’s t test. p>0.05 was considered as not significant and by convention *p<0.05, **p<0.01 and ***p<0.001.

## ACKNOWLEDGEMENTS

The experiments were performed within the France-BioImaging national research (ANR-10-INSB-04, “Investments for the future”), at Montpellier Ressources Imagerie facility (MRI), Montpellier. We thank the engineers on the MRI facility and the members of the team for discussions on the project. This work was supported by the ANR “Chaire d’excellence” CilMitoCyst (ANR-12-CHEX-005 to BD), the Marie Curie career integration grant (CilMitoPatho to BD), the Fondation pour la Recherche Médicale (Partenariat Fondation Schlumberger pour l’Education et la Recherche to BD), the Fondation ARC pour la Recherche sur le Cancer (BD) and the CNRS (BV, CA, BD).

## AUTHORS’ CONTRIBUTION

BD and NT jointly conceived and supervised the project, designed and analyzed the experimental work, assembled the figures and wrote the manuscript. NT technically trained AD and they executed the experimental work. CA helped for cell culture. BV carried out and analyzed the Auxin inducible degron data with the technical help of JM and helped with manuscript editing. VG and YD set up the MicroPoint module on the MRI platform and together with NT optimized it for MT laser ablation.

## DATA AVAILABILITY

The authors declare that the main data supporting the findings of this study are available within the article.

---

**Supplementary movie 1:** Movie from time-lapse microscopy of monopolar Emerald-IFT88 LLC-PK1 cells labelled for tubulin (SiR-Tubulin) showing IFT88 accumulation at minus-end after k-fiber laser ablation (white arrow, ablation site). Movie is displayed at 3 frames per second.

**Supplementary movie 2:** Movies from time-lapse microscopy of monopolar GFP-α-tubulin LLC-PK1 labelled for DNA (Hoechst live, red), to allow for k-fibers detection, in control and IFT88-depleted cells before and after k-fiber ablation (white arrow, ablation site). Movies are displayed at 3 frames per second.

**Supplementary movie 3:** Movies from time-lapse microscopy of monopolar YFP-NuMA LLC-PK1 combined with MT labelling (SiR-Tubulin) in control and IFT88-depleted cells before and after k-fiber laser ablation (arrow, NuMA accumulation at newly generated minus end in control but not in IFT88 depleted condition). Movies are displayed at 3 frames per second.

**Supplementary movie 4:** Movies from time-lapse microscopy of LLC-PK1 GFP-α-tubulin/mCherry-H2B cells showing spindle reorganization and chromosomes alignment in control and IFT88-depleted cells upon nocodazole washout. Movies are displayed at 10 frames per second.

